# EpiPAMPAS: Rapid detection of intra-protein epistasis via parsimonious ancestral state reconstruction and counting mutations

**DOI:** 10.1101/2024.12.13.628430

**Authors:** Fawaz Dabbaghie, Kristina Thedinga, Georgii A Bazykin, Tobias Marschall, Olga Kalinina

**Author notes:** These authors contributed equally (Fawaz Dabbaghie, Kristina Thedinga). Co-last authors (Tobias Marschall, Olga Kalinina).

## Abstract

**Motivation:** An epistatic interaction is a non-linear combination of effects of individual mutations on fitness. This type of interaction is a known driver for evolution, as they alter the organism’s fitness and adaptability. In this work we introduce EpiPAMPAS, a statistical method that is based on multiple sequence alignments (MSA) and detecting mutations in the same direction on a dendrogram instead of a phylogenetic tree using the Sankoff algorithm.

**Results:** We tested EpiPAMPAS on both simulated and real sequencing data. On the simulated data, our method was able to detect the simulated epistatic pairs with very low p-value. In a real-world application, we tested the influenza proteins N1, N2, H1, H3 and HIV-1 envelope protein subtypes A, B and C. We observe that EpiPAMPAS detects fewer interacting pairs than comparable statistical approaches, although the overlap between detected positions is good. Moreover, some of the amino acids from the detected pairs are known to be deleterious for viral fitness.

**Availability:** EpiPAMPAS is available under MIT license at https://github.com/kalininalab/EpiPAMPAS

## 1. Introduction

The first mention of the term epistasis was by William Bateson more than a 100 years ago (3). He used this notion to describe the effect that an allele at one locus had on another allele at another locus. Since then, several definitions have been introduced, but generally, epistasis can be defined as non-additive interaction between loci. On the scale of a single gene or consequently a single protein, two point mutations can be epistatically interacting, if the two mutations can affect each other with respect to their influence on the protein function, which can be detected statistically by observing their mutation pattern during evolution (31).

It has been long noted (37) that a single substitution can depend on the presence or absence of one or more other substitutions in a protein. Later on, several studies have shown that epistatic interactions between mutations are in principle possible (5, 21, 27), and more recently in the context of compound heterozygosity in humans (14). However, as pointed out by (11), the role of epistasis on protein evolution is still not very clear, and some questions remain without a precise answer.

It has been demonstrated that more than half of the missense mutations can be deleterious, that is they reduce the fitness of the organism (19). However, it has been observed that compensatory mutations on other sites in the same gene can help to restore fitness (25, 26, 34). This kind of adaptive epistatic interaction can have an important role in many human diseases for example (16). This mechanism of compensation is known as positive epistasis.

This adaptive evolution can be observed more clearly in viruses that evolve faster than any other organisms and are always under evolutionary pressure to evade human immunity or develop resistance against drugs. For example, many studies on influenza A, and especially on the the viral surface proteins hemagglutinin (HA) and neuraminidase (NA), found high rates of substitutions of the amino acids in these surface proteins (7, 32). Further studies found that epitopes in both HA and NA bind to human antibodies (1, 13, 36), resulting in faster adaptive evolution to evade the immune system.

Looking at the important role that epistasis has, the development of robust statistical models to detect it is important. Several methods based on sequences have been developed to that end, where these methods would look at covariation or mutual information using sequence alignment (10, 17). However, these methods did not take into consideration the phylogenetic relationship between these sequences. Later on, methods that take the phylogenetic information into account have been developed (9, 30).

In particular, a novel statistical method for detection of positive epistasis from phylogenetic data has been developed (18). The method is based on the assumption that under positive epistasis, nonsynonymous substitutions at linked sites follow each other in rapid succession. It involves reconstruction of the phylogenetic tree and of the phylogenetic positions of substitutions. For each pair of sites *i* and *j*, designated as “leading” and “trailing”, they then measure the epistatic statistics *E*(*i, j*) which estimates the time intervals between substitutions occurring at these sites. This statistic is elevated for those site pairs where the time interval between substitutions is, on average, much shorter than that expected on the basis of the overall rate of evolution at this site and the shape of the phylogenetic tree. This approach has been applied to detect epistasis both within (18) and between (24) the two surface proteins of Influenza A, HA and NA, of viral subtypes H1N1 and H3N2. We have established the prevalence of epistasis and determined the pairs of epistatically interacting sites. In particular, those pairs included sites responsible for resistance to oseltamivir - an antiviral drug branded as Tamiflu. Oseltamivir resistance is conferred by the His274Tyr substitution in NA of subtype H1N1 viruses (22), which by itself reduces viral fitness (2). Substitutions Arg222Gln and Val234Met restore viral fitness in vitro (6). It was shown (18) that substitutions at sites 222 and 234 are in epistasis with substitutions at site 274. Out of the 10 most significantly interacting site pairs in NA, six involve site 274. These substitutions are those that can, in combination with His274Tyr, confer oseltamivir resistance. These statistical approaches, although powerful, require reconstruction of the full phylogenetic tree with predicted sequences in all internal nodes, which can be a heavy computational task, depending on the sample size.

In this study, we propose a fast and simple method that relies on hierarchical clustering of sequences instead of full phylogeny reconstruction. Additionally, instead of detecting events that happen faster or slower than expected after each other (as introduced in Kryazhimskiy and Neverov), it detects pairs of mutations that happen more than once independently in this proxy of the phylogenetic tree. We demonstrate the robustness of the method on simulated data and apply it to Influenza A HA and NA sequences for a direct comparison with (18) and HIV-1 envelope protein from subtypes A, B, and C. We demonstrate an agreement of our results with (18) and evaluate the spatial distribution of the epistatically interacting pairs in the protein three-dimensional structures.

## 2. Methods

### 1 Sankoff Algorithm

The proposed method bases the detection of epistasis on a dendrogram, instead of a properly calculated phylogenetic tree, representing the evolutionary relationships between the analyzed samples. The dendrogram is built on multiple sequence alignments (MSA) of amino acid sequences using hierarchical clustering with Ward’s clustering criterion (23, 35). Once the dendrogram has been constructed, for every pair of protein positions, the leaves are labeled with their “genotypes”. Each leaf corresponds to one sample and the labels 0 and 1 represent the two different amino acids observed at each position. To reconstruct the states of the inner nodes of the dendrogram, which correspond to the ancestral states of the protein locations of interest, the Sankoff parsimony (8) is applied to the dendrogram with labeled leaves. The Sankoff parsimony (28, 29) counts the minimal number of evolutionary changes or mutations in a phylogenetic tree – here represented by the dendrogram – to find the most likely ancestral state of a certain locus. In the first step, each node of the phylogenetic tree is assigned a cost vector. This cost vector contains one cell for each possible state of the node storing the minimal evolutionary cost, which is the minimal number of evolutionary changes in the phylogenetic subtree rooted at this node. In the case of protein loci, the possible evolutionary states are 0 and 1, indicating the two different amino acid variants. Initially, the cell in the cost vector corresponding to the evolutionary state observed in each leaf node is set to 0 since there are no evolutionary changes necessary to reach the observed state and all other cells in the cost vector are set to infinity since they are infeasible for the leaf node. The cost vectors of the inner nodes are still unknown at this point, so they are also assigned infinity. Then, starting from the leaf nodes, the cost vectors of all inner nodes up to the root of the phylogenetic tree are calculated according to the following formula:

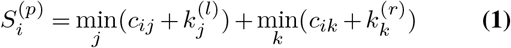

where *i* is the evolutionary state, p denotes the parent node whose cost vector is to be calculated, l and r are the two child nodes of p with already known cost vectors *S*^(*l*)^ and *S*^(*r*)^, and *c*_*ij*_ are entries of the cost matrix *C* containing the costs of evolutionary changes from state *i* to state *j*, which in our case are 0 if *i* = *j* and 1 if *i* ≠ *j*. Thus, the total cost for reaching the states of the leaf nodes in the subtree of node *p* from a state *i* in node *p* is the cost of transitioning from state *i* to the states of the child nodes of *p* plus the minimal cost to reach the states of the leaf nodes from the child nodes.

Supplementary Section 1.1 and Supplementary Figure 1 show an example of a dendrogram with its genotypes, and explains step-by-step how the Sankoff algorithm calculates the states of the inner nodes.

### 2 Dendrogram-based method with mutation direction

We propose a dendrogram-based method to identify epistatic interactions between pairs of protein locations. In this method, a dendrogram is used to account for population structure introduced by evolutionary relationships between the samples of interest. Furthermore, the method takes into account the order in which mutations occur within the dendrogram to distinguish between epistatic and random effects. The underlying idea is that if there are epistatic interactions between two protein locations that are being mutated over time, mutation events at these two locations will most likely not occur independently from each other due to selection pressure. For instance, if two locations *a* and *b* are linked by epistasis and a mutation introducing a variant at location a takes place that decreases the fitness of the organism, it is likely that location *b* will also be mutated, compensating the effect of the variant at location *a*. On the other hand, if locations *a* and *b* both have variants compensating each other, it is less likely that a mutation changing only one of the locations *a* or *b* occurs.

After reconstructing the ancestral evolutionary states of the pair of protein locations that are analyzed for epistasis with the Sankoff parsimony, each of the two protein locations is analyzed for mutation directions with respect to the other location. To this end, mutations in both directions are counted across the whole dendrogram for each of the two locations. If somewhere in the dendrogram the protein location under consideration mutates to the same state (i.e., 0 or 1) as the other location in the pair at the same node in the dendrogram, this is considered a *same direction* mutation, while mutations leaving the pair of protein locations in different evolutionary states are counted towards the *opposite direction* mutations. Pairs of protein locations that are mutated independently are expected to show comparable numbers of *same direction* and *opposite direction* mutations and should thus fit into a binomial distribution with probability of 0.5, while location pairs linked by epistasis do not mutate independently and are expected to deviate from the binomial distribution. Hence, to detect epistasis, we apply two-tailed binomial tests with probability 0.5 on the counts of *same direction* and *opposite direction* mutations obtained from the dendrogram for each of the locations in the pair to obtain p-values.

Supplementary Section 1.2 and Supplementary Figure 2 explain in further detail how this part is done.

### 3 Implementation

EpiPAMPAS is mostly written in R and requires a minimum number of external libraries to run, one preprocessing script written in Python is used to preprocess an MSA into tables that the R modules use.

Figure 1 is an example of an MSA of 5 sequences, it shows how these sequences are preprocessed by the Python module, producing a VCF and a simple table of all possible pairs combinations of variant loci. The VCF-like table is then turned into a matrix with rows as samples and columns as possible variant positions with values as the “genotype” in the R Module. For each pair of positions, only the samples that contain that pair of positions in the matrix are kept, i.e. if there was a gap at that position like in sequence 1 position 7, then the value will be NA in the matrix. The matrix is then used to build the dendrogram using hierarchical clustering, and the most likely genotype is reconstructed (the genotype that requires the least number of mutations) using Sankoff parsimony.

**Fig. 1.**
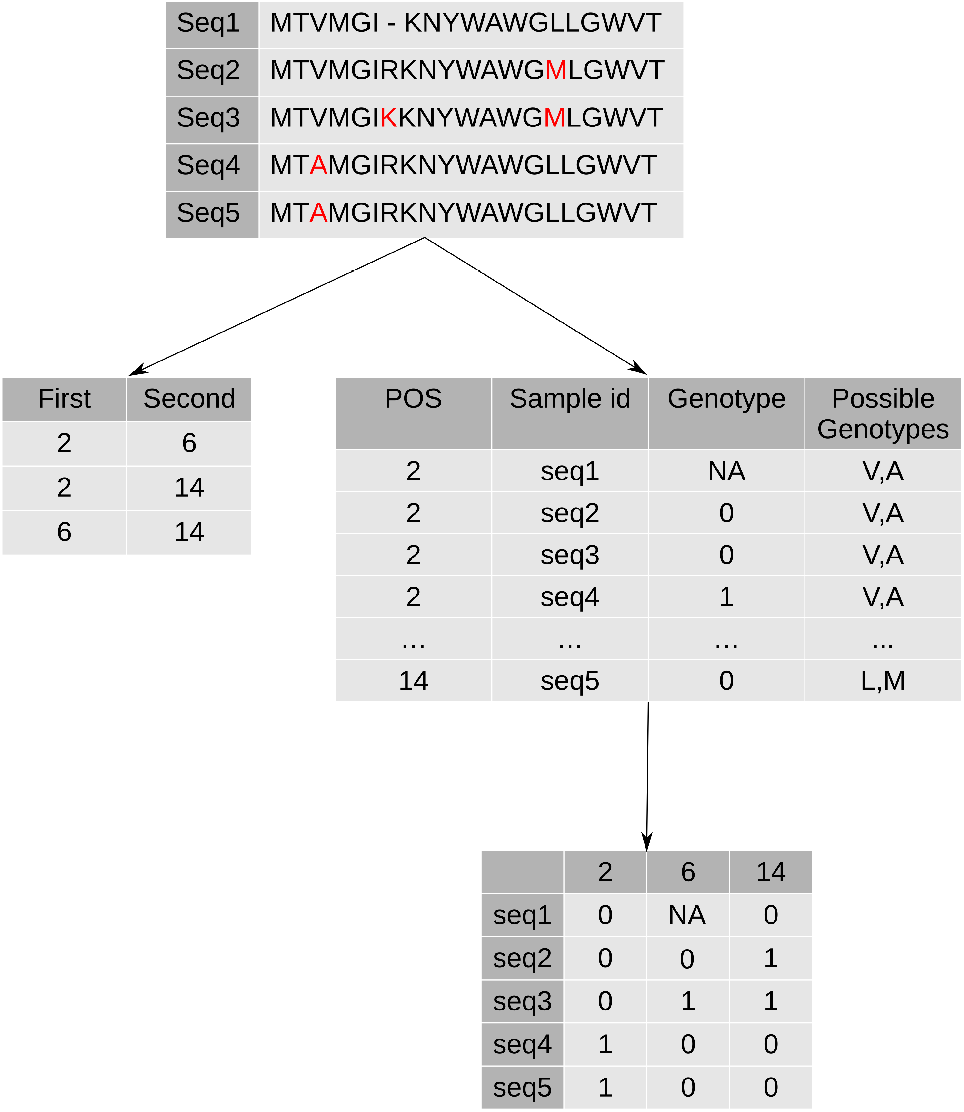
Preprocessing of MSA: An example of an MSA with 5 sequences and 3 variable positions, the middle table to the left is the 3 possible pairs of these 3 positions in this MSA, the middle right table is the VCF table with the information related to each sample (0 indexing is used here), the variant position and the variant value. The last table is the matrix representing the VCF-style table that is used to build the dendrogram

## 3. Data

### 1 Simulated Data

To test that our dendrogram-based algorithm does actually detect a pair of mutations that independently happen in the same direction, simulated dendrograms were used. In the first step, balanced dendrograms are constructed from n samples, where each node has exactly two child nodes. We then mark the nodes with the same genotype “0” which denotes that for a pair of positions, both positions have the wildtype genotype. We start introducing same direction mutations where both positions mutate to have the same genotype, and opposite direction mutations where they have different genotypes. This introduction is done with different probabilities, where *p*_*same*_ is the probability of introducing same direction mutations and *p*_*opposite*_ is the probability for introducing opposite mutations. In this example we vary *p*_*same*_ between 0 and 0.5 with 0.05 steps, and poppositeis defined with this equation ,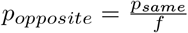 where *f* is a factor between 1 and 4.

Mutations are introduced starting from the root node of the tree, where each node is mutated with probability *p*_*same*_ + *p*_*opposite*_, and if the mutation is introduced at some node v, the genotype is propagated through the subtree rooted at v. This mutation procedure is repeated 100 times for each parameter combination and each number of samples (10, 50, 100, 500) to get a representative pool of results. (See Results section for the results of the simulated experiment).

### 2 Real Data

Two datasets were used for testing our method. First, we used the same dataset of Influenza A hemagglutinins and neuraminidases of subtypes H1, H3 and N1, N2, respectively, as in the (18) study in order to enable a direct comparison with their results. The second dataset is the HIV-1 envelope gp160 glycoprotein (Env) sequences from HIV-1 subtypes A, B, and C, taken from the Los Alamos National Laboratory (LANL) HIV Sequence Database (20).

To investigate the potential functional impact of the detected epistatic interactions, we investigated their location in the corresponding protein three-dimensional structures. For each protein, a structure from the Protein Data Bank (4) was chosen, such that for the Influenza proteins we used the same structures as described in (18), and for the HIV-1 envelope we searched for the most complete structures of gp120 and gp41 (the end products in the env gene) from the corresponding subtypes using StructMAn (12) (Tables 1 and 2). Bonferroni correction was used on the p-values calculated with our method to correct for multiple testing in case when more than one mutation pair can be observed in the same pair of positions (e.g., when one alignment position contains three or more different amino acids). In order to calculate the pairwise distance between the detected pairs, the sequences were mapped onto the structures, and the pairwise distances between the nearest atoms in the corresponding amino acids were calculated. Afterwards, the Wilcoxon signed-rank test was calculated between the pairs we identified as possibly epistatically interacting and all possible pairwise distances.

**Table 1.** Influenza A Dataset: Real-world dataset used in this study on Influenza A hemagglutinin and neuraminidase sequences. Bonferroni multiple tests correction applied.

| Protein | H1 | H3 | N1 | N2 |
| --- | --- | --- | --- | --- |
| # Sequences | 1219 | 2149 | 1836 | 2339 |
| # Pairs p-value <0.1 | 7 | 33 | 19 | 18 |
| # Pairs p-value <0.05 | 3 | 23 | 1 | 7 |
| # Pairs p-value <0.01 | 1 | 5 | 0 | 3 |
| Structure PDB ID | 1RUZ | 2VIU | 3BEQ | 1NN2 |

**Table 2.**
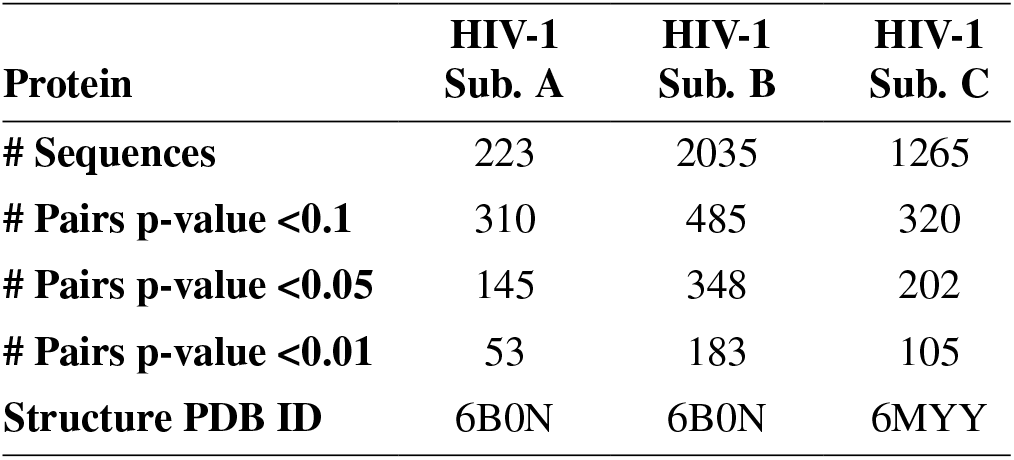
HIV Dataset: Real-world dataset used in this study on HIV-1 Env sequences. Bonferroni multiple tests correction applied.

## 4. Results

### 1 Simulated Data

Testing on simulated data confirms the validity of our approach (Figure 2): the higher the value for *p*_*same*_ (which indicates more mutations in the same direction, but only if *p*_*same*_ is larger than *p*_*oppposite*_), the lower the p-values are detected by our method. Moreover, larger values of *f* (which result in smaller values for *p*_*opposite*_ compared to *p*_*same*_) also result in more significant p-values. We can also see that the number of samples has an effect on the p-value reported by our method: the bigger the sample size is, the more significant the reported p-value.

**Fig. 2.**
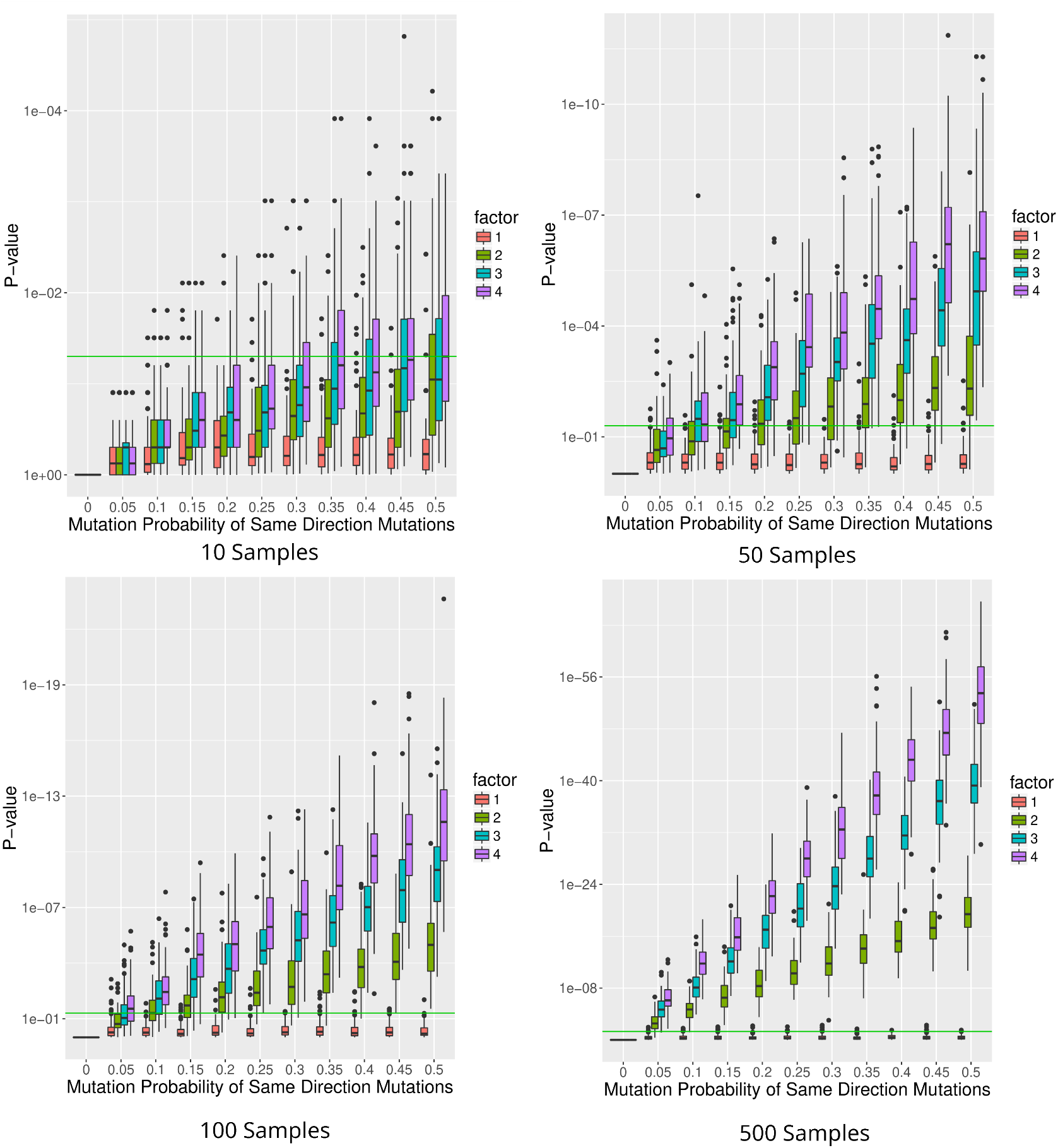
Simulated data: Boxplots for the simulated trees. Different sample sizes were used for this test. The x-axis represented the different *p*_*same*_ probabilities, the different colors for the boxes represent the different *f* values, and the y-axis represents the p-value measured by our method. The green line indicates the 0.05 p-value.

### 2 Application to Viral Proteins

For the real data, first, we ran our algorithm on the Influenza A dataset of proteins H1, H3, N1, and N2 from (18). After filtering using a p-value threshold of 0.05, we were left with few potentially epistatically interacting pairs (Table 1), so we opted to use a more relaxed cutoff of 0.1. Notably, (18) predicted many more epistatically interacting pairs, in line with their mentioned higher false discovery rate (FDR). Nevertheless, the results of both methods agree very well (Figure 5, right). Both methods detect the same sets of amino acid positions, however the predicted interacting pairs (Figure 5, left) are largely different. Only for H1, we detect three interacting pairs, which were all discovered by (18).

We compared the distance distribution of detected potentially epistatically interacting pairs with the background distance distribution of all amino acids in the corresponding protein three-dimensional structures using Wilcoxon one-sided signed-rank test (Tables 3 and 4). We see that in influenza, the proteins H1, H3, and N1 had a significant p-value but only in H3 the distance was significantly smaller (Figure 3). However, with H1 and N1, the number of pairs was much smaller compared to H3. For HIV1 we see that the p-value is significant for subtypes B and C, but looking at Figure 3, the average distances are not that different. Moreover, the corresponding AUC values are low, indicating that this significance probably is simply due to the higher number of detected pairs in these datasets.

**Table 3.** Wilcoxon one-sided test in influenza comparing distances between potentially epistatically interacting pairs detected using our method for p-value threshold of 0.1 with all pairwise distances in the structure.

| Protein | H1 | H3 | N1 | N2 |
| --- | --- | --- | --- | --- |
| Test statistic W | 35391.5 | 908535.5 | 6539.5 | 6885 |
| P-value | 0.04 | 0.005 | 0.051 | 0.466 |
| AUC | 0.79 | 0.35 | 0.97 | 0.48 |

**Table 4.** Wilcoxon one-sided test in HIV: Comparing distances between potentially epistatically interacting pairs detected using our method filtered against distances between all amino acid pairs for p-value <0.05.

| Protein | HIV-1<br>Sub.<br>A | HIV-1<br>Sub.<br>A | HIV-1<br>Sub.<br>A |
| --- | --- | --- | --- |
| Test statistic W | 309836.5 | 1005287 | 642826 |
| P-value | 0.308 | 0.0002 | 0.088 |
| AUC | 0.52 | 0.6 | 0.54 |

**Fig. 3.**
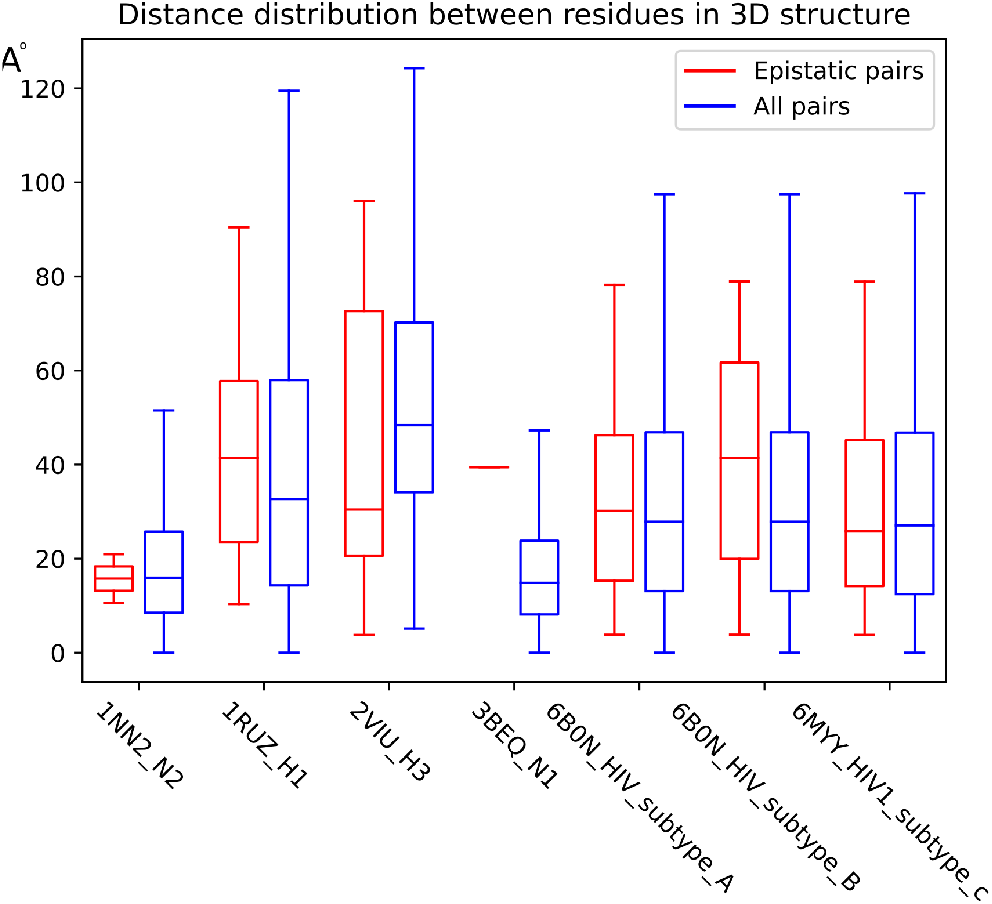
3D Structure distances: Boxplot with distance distribution of all pairs in a 3D structure (in blue) and only the epistatic interacting pairs that we detected (in red).

In the HIV1 dataset, looking at the overlap between the pairs for each subtype, we found that between subtypes A and B only 2 pairs were the same, between A and C also only 2 pairs, and between A and C only 4 pairs overlap; this is small number compared to the actual number of pairs for each subtype which is 145, 348, and 202 for subtypes A, B, and C respectively. This points to the evolutionary diversity between the subtypes that has been seen before, which adds to the challenges of effectively controlling the virus (33).

In influenza, some mutations have been associated with resistance to oseltamivir, e.g., mutation in position 274 the mutation from Histidine to Tyrosine (H274T) has been shown to give the virus resistance to oseltamivir drug (22). However, in (2), it was demonstrated that this mutation is deleterious in the absence of the drug. Moreover, mutations at position 222 between Arginine and Glutamine (R222Q) and at position 234 between Histidine and Tyrosine (H234Y) regain the viral resistance against the drug (6). Both R222Q and H234Y were detected by our method with significant p-values.

Comparing our detected positions in the HIV sequences with a list of biologically relevant residues, such as those in the CD4 binding site or antigenic epitopes taken from (15), looking at Table 5 we see that the fraction of amino acid residues in biologically important regions from (Hake and Pfeifer 2017) among detected potentially epistatically interacting positions is around 39%, 43%, and 42% for HIV-1 subtypes A, B, and C respectively. For all amino acids in Env, the fraction of the ones in biologically relevant regions is around 32%.

**Table 5.**
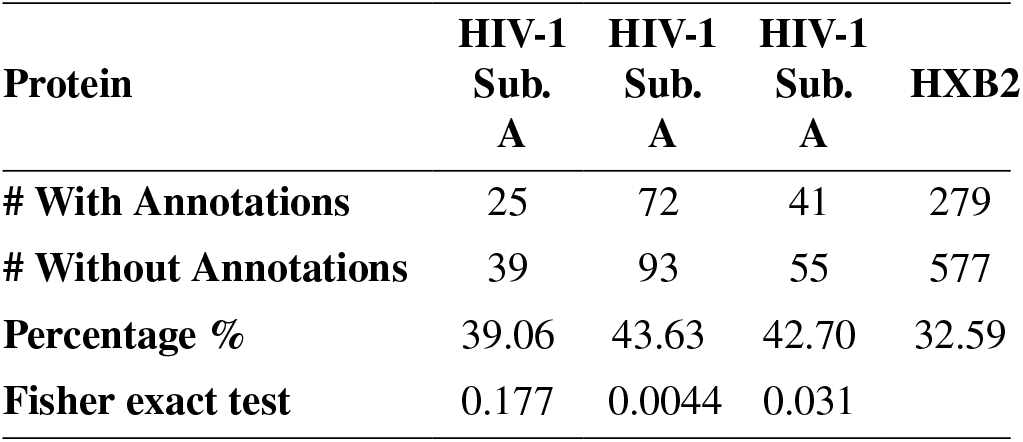
HIV Positions Annotations: Using of annotated positions in the HIV1 envelope protein taken from (15), we looked at how many of the positions we detected had annotation compared to the reference protein, and ran a Fisher exact test to see if the inflation in the number of annotated positions to not annotated positions is more significant in the positions we detected.

To see if this increase of percentage is significant, we ran a Fisher exact test and we see that for HIV-1 subtypes B and C, the p-value is smaller than 0.05. The structural classes (e.g. interactions with other proteins or small ligands, core and surface residues) of all detected positions were evaluated using StructMAn and compared to the background distribution of structural classes for all Env amino acids (Figure 4). Potential epistatically interacting positions from all three subtypes follow a similar trend with a slight depletion from the core compared to the background. Interestingly, there is no enrichment on the protein-protein interaction interfaces, which is in a slight contrast with the previous observation, since most biologically relevant residues from (15) signify an interaction with another protein, and epistatically interacting residues are enriched among them. Hence, epistatically interacting residues must be less common at the interaction interfaces between the Env subunits in the trimer.

**Fig. 4.**
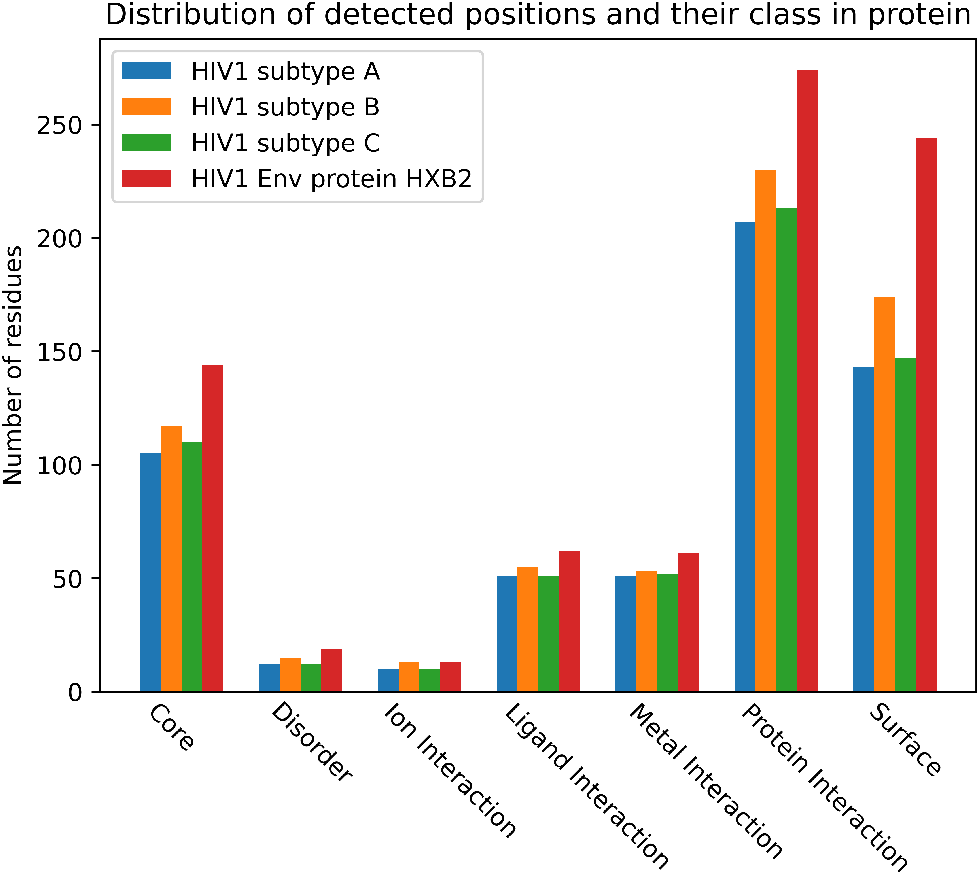
Class distribution in detected residue: Bar plot with distribution of number of detected residues as potential epistatic pairs against the reference sequence HXB2.

**Fig. 5.**
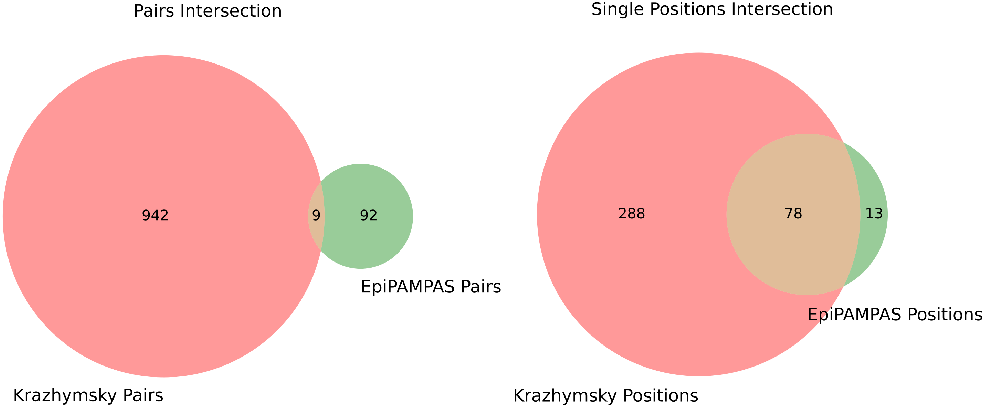
Results Venn Diagram: this plot is showing the intersection between the individual positions detected between our method and (18) method, and the intersection of the detected pairs. We see that the majority of our positions were also detected in their results. However, when it comes to pairs, the intersection is much smaller, indicating that the pairs we detected are different from their pairs. The results represented here are filtered for a p-value smaller than 0.05 except for N1, where we used 0.1 because there was only one pair detected with a p-value cutoff of 0.05.

## 5. Conclusion

In this paper, we presented EpiPAMPAS, a new statistical method for detecting epistatic interactions between mutations. Our method is based on counting same-direction mutations on a dendrogram, in contrast to other tree-based methods that require building a complete phylogenetic tree. We tested our method on both simulated and viral protein data. We showed that for simulated data, EpiPAMPAS performed well. For viral protein data, we compared our method to another tree-based method. EpiPAMPAS many positions that were also identified by (18) , but reported fewer positions in total. However, compared to (18), the interactions between the positions are detected differently. Some of the mutations we detected are known to be deleterious from the literature. From the protein structure perspective, we did not find a clear signal that the pairs detected are closer to each other in the 3D protein structure compared to all possible pairs of mutations. One reason for that is that our statistical definition of epistasis does not reflect well the actual fitness-based definition. Another possibility, which we consider more likely, is that the sequence data on viral proteins, despite our best effort to collect as much of it as possible, is still insufficient to detect epistasis. Two arguments in favor of this view can be, first, a good performance for simulated data, and second, the fact that for some datasets we detect as few as one pair for p < 0.05. Another explanation could be that the way we simulated the data does not reflect the real mechanisms on how compensation due to epistasis happens, and that our simulation procedure is positively biased towards our methods.

