## Supplementary Materials for "EpiPAMPAS: Rapid detection of intra-protein epistasis via parsimonious ancestral state reconstruction and counting mutations"

[1]Institute for Medical Biometry and Bioinformatics, Medical Faculty and University Hospital Düsseldorf, Heinrich Heine University Düsseldorf, 40225 Düsseldorf, Germany
[2]Center for Digital Medicine, Heinrich Heine University, 40225 Düsseldorf, Germany
[3]Helmholtz Institute for Pharmaceutical Research Saarland (HIPS), Helmholtz Center for Infection Research (HZI), Saarbrücken, Germany
[4]Max Planck Institute for Molecular Genetics, Bioinformatics Department, Ihnestr. 63-73, 14195, Berlin.
[5]Department of Biology, Emory University, Atlanta, Georgia 30322, USA
[6]Drug Bioinformatics, Medical Faculty, Saarland University, 66421 Homburg, Germany.
[7]Center for Bioinformatics, Saarland University, 66123 Saarbrücken, Germany
[*]These authors contributed equally. [†]Co-last authors

December 13, 2024

### 1 More on Sankoff and Mutation Counting

#### 1.1 Sankoff Parsimony Method and Example

In Supplementary Figure 1, we see an example of a dendrogram with 13 samples (leaf nodes), and the cost vector for each node in the tree is calculated step-by-step in the figure. Here, our cost vector $S^{(p)}$ has two possible states representing the genotype. If more than two genotypes are available for that position, we subset our samples and take all combinations of two genotypes for a pair of positions and calculate a tree based on this subsample.
We label our two genotypes as “o” and “x”, where the left cell in the cost vector represents the genotype “o” and the right cell represents the “x” genotype. When initializing the tree with the cost vector of the leaf node, the genotype of that leaf node gets the value 0, and the other genotype is assigned $\infty$. For example, in the dendrogram in the Supplementary Figure 1, looking at the left pair of leaf-nodes, the first one has the genotype “x” and is initialized with $[\infty, 0]$, and the second leaf node has the genotype “o” and is initialized with $[0, \infty]$. Once the leaf nodes are labeled, we can start calculating the values for each parent node using Equation 1, because our cost value is of size two and we only have two genotypes, we can expand that equation and calculate the values for each node as follows:

$$\begin{aligned} node[o] &= min(lcn[o], lcn[x] + 1) + min(rcn[o], rcn[x] + 1) \\ node[x] &= min(lcn[o] + 1, lcn[x]) + min(rcn[o] + 1, rcn[x]) \end{aligned} \quad (1)$$

where $node[o]$ is the value for genotype “o” in the node, and $node[x]$ the value for genotype “x”; $lcn$ means “left-child node”, and $rcn$ means “right-child node”. For example, looking at step 4 in the figure, in the middle we have the calculated vector $[1, 2]$, under that node, we have two leaf nodes ($[0, 2]$ and $[1, 1]$), the first value “1” was calculated with $(min(0, 2 + 1) + min(1, 1 + 1) = 0 + 1 = 1)$, and the second value “2” was calculated with $(min(0 + 1, 2) + min(1 + 1, 1) = 1 + 1 = 2)$. Same goes for the root node $[4, 4]$, this was calculated using the two child nodes ($[2, 1]$ and $[2, 4]$) and the first

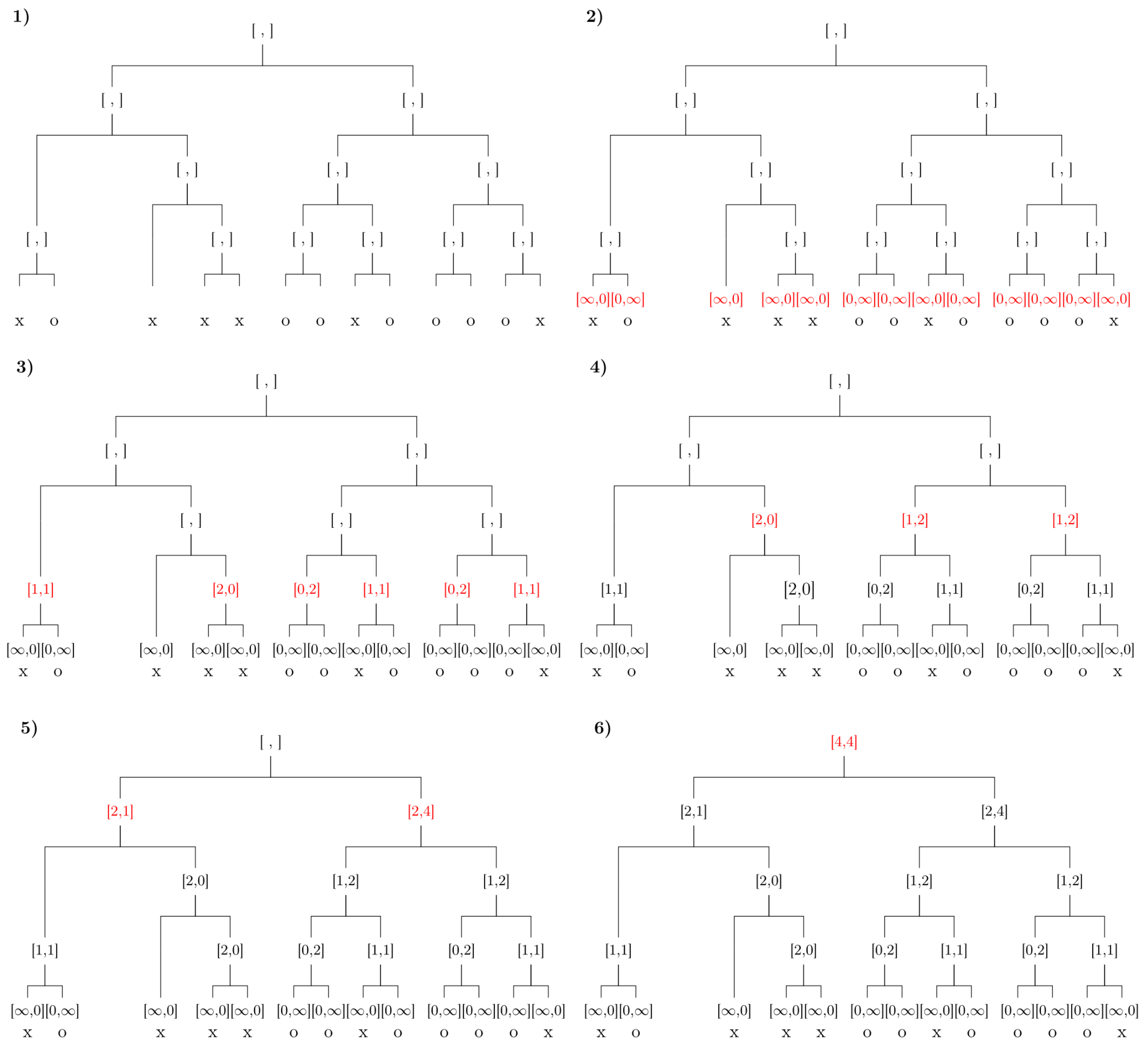


**Supplementary Figure** 1: Example on how Sankoff algorithm works

value calculated with $(min(2, 1 + 1) + min(2, 4 + 1) = 2 + 2 = 4)$ and the second value calculated with $(min(2 + 1, 1) + min(2 + 1, 4) = 1 + 3 = 4)$.

### 1.2 Mutation counting

Once we finished with the Sankoff algorithm and constructed the state of the inner nodes, i.e., reconstruct the most likely genotype of the inner nodes. We traverse the tree starting from the root and count the number of same and opposite direction mutations. We do two counting runs. First, we consider the first position in the protein position-pair as constant and count the mutation direction in the second position (as seen in Supplementary Figure 2. Second, we consider the second position in the protein position-pair as constant and count the the mutation direction of the first position.
Supplementary Figure 2 also shows the two tables of the two possible counts, where same direction mutations are the sum $(b_2 + c_2)$ and opposite direction mutations are the sum $(a_2 + d_2)$.
In our method, we expect that variants/positions that do not have an epistatic interaction would fit a binomial distribution with a probability of 50% and the ones with an epistatic interaction would deviate from this distribution. EpiPAMPAS offers the user to do a two-sided or one-sided binomial distribution, where a one-sided can be then used to test if the count of one direction occurs more often than the other direction.

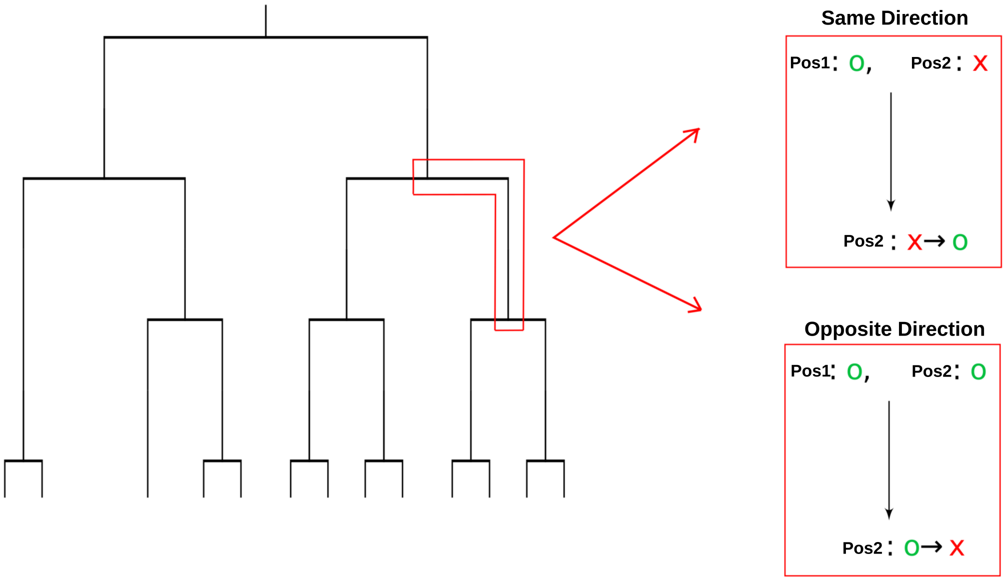


| | **Pos1** : $o$ | **Pos1** : $x$ |
|---|---|---|
| **Pos2**: $o \to x$ | $a_1$ | $b_1$ |
| **Pos2**: $x \to o$ | $c_1$ | $d_1$ |

| | **Pos2**: $o$ | **Pos2**: $x$ |
|---|---|---|
| **Pos1** : $o \to x$ | $a_2$ | $b_2$ |
| **Pos1** : $x \to o$ | $c_2$ | $d_2$ |

**Supplementary Figure** 2: The tree shows the same direction and opposite direction mutations of the second position (Pos2) while keeping (Pos1) constant. In red, is one event in the inner tree we are looking at, where we count 1 for the same direction mutation (top red box) if the mutation in the second position follows the first position and mutates to the same genotype, and count 1 for opposite direction mutations (bottom red box) if the mutation results in different genotypes.

### 2 Supplementary Result Figures

Supplementary Figure 3 showcases the different Venn diagrams for the comparison between EpiPAMPAS results and the results from [1]. Each row in the figure represent a different protein, the venn diagrams on the left are for the intersection of the positions detected by both methods, and the ones on the right show the intersection of the pairs detected.

Supplementary Figure 4 shows scatter plots between the 1D and the 3D distance of the pairs detected in the proteins.

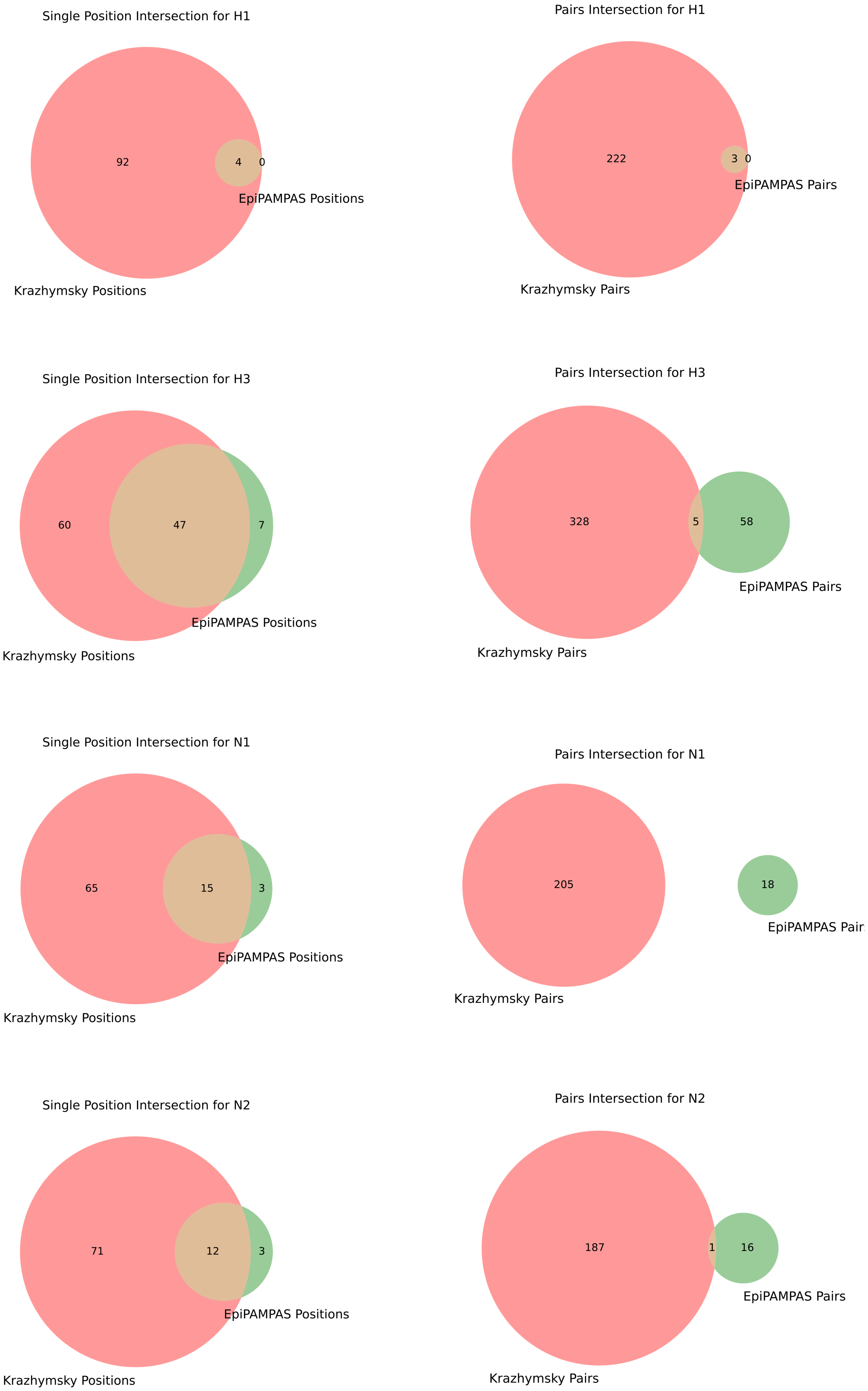


**Supplementary Figure** 3: Intersection between the positions and the pairs of positions EpiPAMPAS detected and the method from (Kryazhimskiy et al. 2011) for each of the viral proteins H1, H3, N1, and N2. We can see that the overlap of positions detected is big. However, the intersection when it comes to the pairs of interacting positions detected is rather small between the two methods.

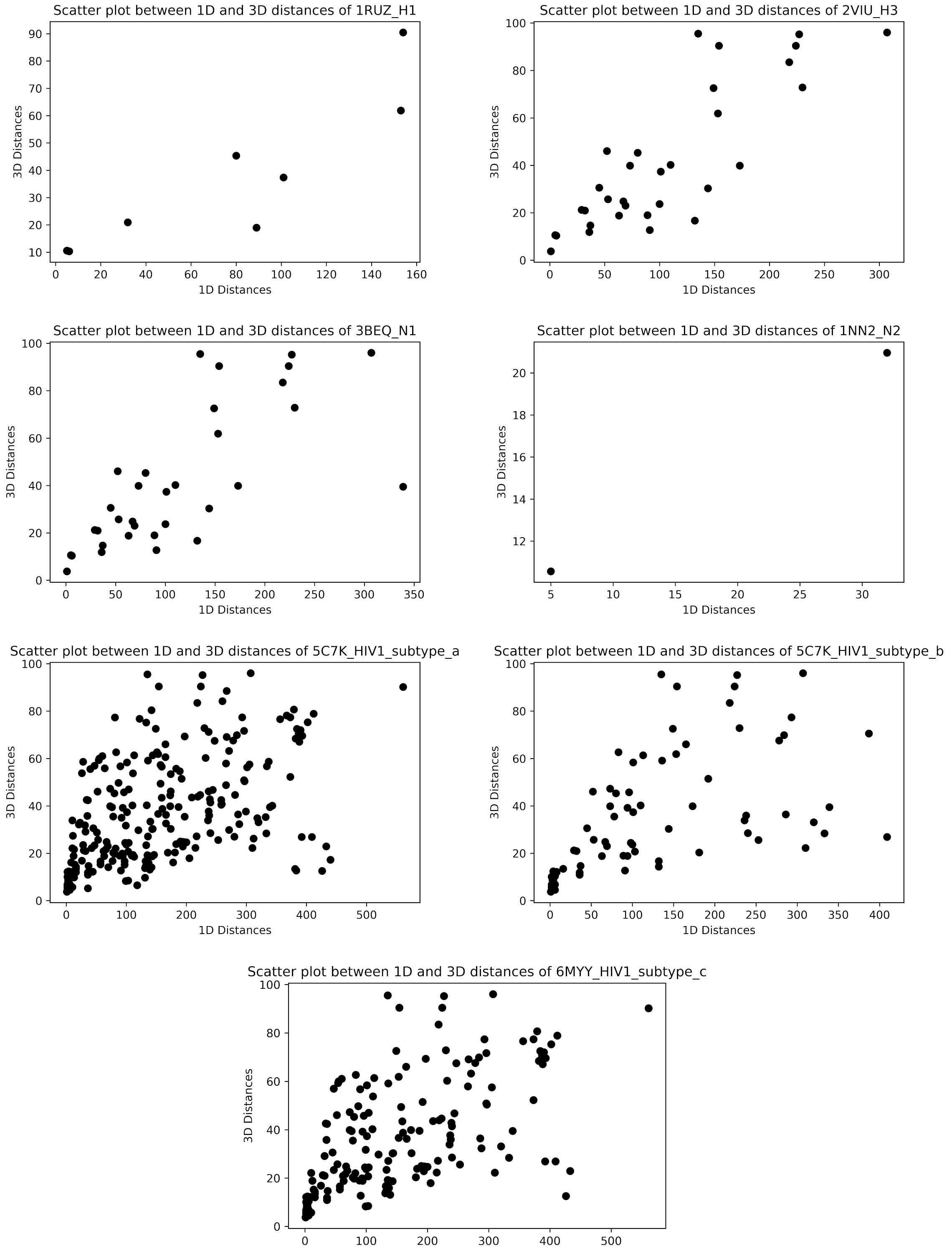


**Supplementary Figure** 4: Scatter plot of the 1D vs 3D distance of the pairs detected with Epi-PAMPAS for H1, H3, N1, N2, HIV1 subtype a, HIV1 subtype b, and HIV1 subtype c using the structures 1RUZ, 2VIU, 3BEQ, 1NN2, 5C7K, 5C7K, and 6MYY respectively. We see that there is a trend where the longer the 1D distance, the longer the 3D distance. However, we would expect more of a trend where the 3D distance is smaller indicating that the pairs detected have some interaction in the 3D structure.
